# Loss of *O*-linked protein glycosylation in *Burkholderia cenocepacia* impairs biofilm formation and siderophore production via alteration of quorum sensing regulation

**DOI:** 10.1101/757575

**Authors:** Cameron C. Oppy, Leila Jebeli, Miku Kuba, Clare V. Oates, Richard Strugnell, Laura E. Edgington-Mitchell, Miguel A. Valvano, Elizabeth L. Hartland, Hayley J. Newton, Nichollas E. Scott

**Author notes:** These authors contributed equally (Cameron C. Oppy, Leila Jebeli).

## Abstract

*O*-linked protein glycosylation is a conserved feature of the *Burkholderia* genus. For *Burkholderia cenocepacia*, the addition of the trisaccharide β-Gal-(1,3)-α-GalNAc-(1,3)-β-GalNAc to membrane exported proteins is required for virulence and resistance to environmental stress. However, the underlying causes of the defects observed in the absence of glycosylation are unclear. This study demonstrates that the global *B. cenocepacia* proteome undergoes dramatic changes consistent with alterations in global transcriptional regulation in the absence of glycosylation. Using luciferase reporter assays and DNA cross-linking analysis, we confirm the repression of the master quorum sensing regulon CepR/I in response to the loss of glycosylation, which leads to the abolition of biofilm formation, defects in siderophore production, and reduced virulence. The abundance of most of the known glycosylated proteins did not significantly change in the glycosylation-defective mutants except for BCAL1086 and BCAL2974, which were found in reduced amount, suggesting they could be degraded. However, the loss of these two proteins was not responsible for driving the proteomic alterations, as well as for reduced virulence and siderophore production. Together, our results show that loss of glycosylation in *B. cenocepacia* results in a global cell reprogramming via alteration of the CepR/I regulon, which cannot be explained by the abundance changes in known *B. cenocepacia* glycoproteins.

**IMPORTANCE:** Protein glycosylation is increasingly recognised as a common protein modification in bacterial species. Despite this commonality our understanding of the role of most glycosylation systems in bacterial physiology and pathogenesis is incomplete. In this work, we investigated the effect of the disruption of *O*-linked glycosylation in the opportunistic pathogen *Burkholderia cenocepacia* using a combination of proteomic, molecular and phenotypic assays. We find that in contrast to recent findings on the *N*-linked glycosylation systems of *Campylobacter jejuni, O*-linked glycosylation does not appear to play a role in proteome stabilization of most glycoproteins. Our results reveal that virulence attenuation observed within glycosylation-null *B. cenocepacia* strains are consistent with alteration of the master virulence regulator CepR. The repression of CepR transcription and its associated phenotypes support a model in which the virulence defects observed in glycosylation-null strains are at least in part due to transcriptional alteration and not the direct result of the loss of glycosylation *per-se*. This research unravels the pleotropic effects of *O*-linked glycosylation in *B. cenocepacia,* demonstrating that its loss does not simply affect the stability of the glycoproteome, but also interferes with transcription and the broader proteome.

## Introduction

The *Burkholderia cepacia complex* (Bcc) includes diverse and ubiquitous, phylogenetically related Gram-negative species (1). To-date, 20 Bcc species have been identified (1–3), but the commonality of Bcc in the environment (2, 3) and their recognition as opportunistic pathogens (4–6) continually drives the identification of new Bcc members. Within clinical settings Bcc can lead to fatal infections (7, 8) which are both challenging to control with antibiotic therapies (9) and can be spread by patient-to-patient transmission (10, 11). This is especially problematic for Bcc infections within cystic fibrosis (CF) patients, where Bcc infections result in accelerated loss of lung function (12) as well as increased morbidity and mortality compared to other infectious agents (13, 14). *B. cenocepacia* is one of the most common Bcc species isolated from CF patients across the globe (15–18) and is generally associated with more fulminant disease leading to higher mortality than observed with other Bcc species (19). One of the most serious clinical outcomes from *B. cenocepacia* infections in CF patients is a condition known as ‘cepacia syndrome’, an unrelenting necrotizing pneumonia that rapidly leads to respiratory failure, bacteraemia and death (20). Although interventions with antimicrobial therapies can stop or even reverse cepacia syndrome (20), the intrinsic resistance of Bcc to multiple classes of antibiotic (21–23) and their propensity to form biofilms (24) makes treatment success variable at best (9). It is therefore essential to better understand factors involved in *B. cenocepacia* virulence in order to improve clinical outcomes.

The ability to form biofilms is associated with bacterial persistence and the failure of antimicrobial treatments in a range of pathogens (25). Bcc members, including *B. cenocepacia,* produce biofilms on abiotic (26, 27) and biotic surfaces (28). However, *B. cenocepacia* in the CF lung do not appear to form true biofilms, but instead are observed extracellularly as small clusters surrounded by mucus, or in phagocytic cells in the submucosal tissue (29, 30). Increased biofilm production is associated with bacterial persistence in CF patients (31) and mutations selected for during chronic infections in CF patients mirror those observed during biofilm directed evolution experiments (32). The ability to form biofilms in Bcc, as well as the expression of multiple virulence factors, is controlled by numerous quorum sensing (QS) systems (33). A key class of QS systems associated with Bcc virulence are based on homoserine lactones (HSL) (24). Across the Bcc, some HSL QS systems are variable or lineage-specific, such as cciR/I and CepR2 (34, 35), while others are highly conserved in all members. One such highly conserved HSL QS system is the CepR/I regulon (36, 37), which generates N-octanoylhomoserine lactone (C8-HSL) using the HSL synthase CepI that in turns activates the transcriptional regulator CepR (37, 38). CepR is a major regulator of biofilm formation (39), and disruption of CepR/I attenuate Bcc virulence in several models (40, 41) and reduces disease severity (40, 42). The importance of the CepR/I QS system in Bcc virulence stems from its broad regulatory profile affecting multiple virulence-associated genes (43–45) such as those encoding the secreted zinc metalloproteases ZmpA (46) and ZmpB (47), siderophore production (39, 48) and the key mediator of <u>B</u>iofilm formation <u>p</u>rotein A (BapA) (45).

Glycosylation is increasingly recognised as a common modification in bacterial systems (49–56). Many glycosylation systems are conserved across bacterial genera (57, 58) and phyla (59, 60), suggesting glycosylation is critical for optimal proteome functionality. Disruption of glycosylation pathways in several species results in reduced fitness compared to glycosylation competent strains (52–56). However, the underlying cause of the reduction in fitness remains poorly defined (61, 62). Only recently have mechanistic insights emerged on how the loss of glycosylation affects bacterial physiology and pathogenesis. In *Campylobacter jejuni*, the loss of glycosylation results in decreased stability of the majority of known glycoproteins, which in turn affects virulence (63, 64). These data support a model whereby bacterial *N*-linked glycosylation contributes to protein stability, but it is unclear whether other glycosylation systems, such as *O*-linked glycosylation, have evolved to stabilize glycosylated proteins.

Previously, we reported *B. cenocepacia* possesses an *O*-linked glycosylation system responsible for the modification of at least 23 proteins with a trisaccharide glycan using the enzyme PglL (BCAL0960) (56). Building on this work we recently identified the biosynthetic locus, the *<u>O</u>*-<u>g</u>lycosylation <u>c</u>luster (OGC, BCAL3114 to BCAL3118) responsible for the generation of the *O*-linked glycan, established the *O*-linked glycan structure as β-Gal-(1,3)-α-GalNAc-(1,3)-β-GalNAc, and demonstrated that glycosylation was required for optimal bacterial fitness and resistance to clearance in *Galleria mellonella* infection models (65). Although these studies have demonstrated a link between glycosylation and virulence, the mechanism remains unclear. Using quantitative proteomic approaches, we sought to understand the proteome changes resulting from the loss of *O*-linked glycosylation in *B. cenocepacia*. We demonstrated that loss of glycosylation in *B. cenocepacia* resulted in global proteome alterations beyond the known glycoproteome, which are associated with widespread alterations in transcriptional regulation. We discovered that the HSL QS system CepR/I is repressed in glycosylation-defective mutants, which leads to defective biofilm formation and reduced siderophore production. In contrast to the loss of glycosylation in *C. jejuni,* we also demonstrate that only a few glycoproteins are reduced in abundance in the absence of glycosylation, but they are not responsible for the glycosylation-null phenotypes. Together, our data indicate that the roles of glycosylation in *B. cenocepacia* extend beyond protein stabilisation, and loss of *O*-linked glycosylation in *B. cenocepacia* causes dramatic physiological changes by altering the global regulatory CepR/I QS system.

## Methods

### Bacterial strains and growth conditions

Strains and plasmids used in this study are listed in Supplementary Table 1 and 2, respectively. Strains of *Escherichia coli* and *B. cenocepacia* were grown at 37°C in Luria-Bertani (LB) medium. When required, antibiotics were added to a final concentration of: 50 μg/ml trimethoprim for *E. coli* and 100 μg/ml for *B. cenocepacia*, 20 μg/ml tetracycline for *E. coli* and 150 μg/ml for *B. cenocepacia* and 40 μg/ml kanamycin for *E. coli*. Ampicillin was used at 100 μg/ml and polymyxin B at 25 µg/ml for triparental mating to select against donor and helper *E. coli* strains. Antibiotics were purchased from Thermo Fisher Scientific while all other chemicals unless otherwise stated were provided by Sigma-Aldrich.

### Recombinant DNA methods

Oligonucleotides used in this study are listed in Supplementary Table 3. DNA ligations, restriction endonuclease digestions, and agarose gel electrophoresis were performed using standard molecular biology techniques (66) with Gibson assembly undertaken according to published protocols (67). All restriction enzymes, T4 DNA ligase and Gibson master mix were used as recommended by the manufacturer (New England Biolabs). *E. coli* pir2 and DH5α cells were transformed using heat shock-based transformation. PCR amplifications were carried out using either Phusion DNA (Thermo Fisher Scientific) or Pfu ultra II (Agilent) polymerases were used according to the manufacturer recommendations with the addition of 2.5% DMSO for the amplification of *B. cenocepacia* DNA due to its high GC content. DNA isolation, PCR recoveries and restriction digest purifications were performed using the genomic DNA clean-up kit (Zmyo research, CA) or Wizard SV gel and PCR clean-up system (Promega). Colony and screening PCRs were performed using GOTAQ Taq polymerase (Qiagen) supplemented with 10% DMSO when screening *B. cenocepacia*. DNA sequencing was undertaken at the Australian Genome Research Facility (Melbourne, Australia).

### Construction of unmarked deletion mutants, endogenous tagged BCAL1086 and complementation with *pglL-his*_10_

Deletions and endogenous tagging of BCAL1086 were undertaken using the approach of Flannagan *et al.* for the construction of unmarked, non-polar deletions in *B. cenocepacia* K56-2 (68). Chromosomal complements of *pglL* were generated by introducing *pglL-his*_10_ under control of the *B. cenocepacia* S7 promoter (*P*_S7_) or the native pglL promoter (Ppgl, 660bp upstream of PglL) inserted into *amrAB* using the pMH447 (23) derivative plasmids (Supplementary Table 2) according to the protocol of Aubert *et al* (69).

### Protein manipulation and immunoblotting

Bacterial whole-cell lysates were prepared from overnight LB cultures of *B. cenocepacia* strains. 1ml of bacteria at an OD_600_ of 1.0 were pelleted, then resuspended in 4% sodium dodecyl sulfate (SDS), 100 mM Tris pH 8.0, 20 mM dithiothreitol (DTT) and boiled at 95°C with shaking at 2000 rpm for 10 minutes. Samples were then mixed with Laemmli loading buffer [24.8 mM Tris, 10 mM glycerol, 0.5% (w/v) SDS, 3.6 mM β-mercaptoethanol and 0.001% (w/v) of bromophenol blue (pH 6.8) final concentration] and heated for a further 5 minutes at 95°C. Lysates were then subjected to SDS PAGE using pre-cast 4-12% gels (Invitrogen) and transferred to nitrocellulose membranes. Membranes were blocked for 1 hour in 5% skim milk in TBS-T (20 mM Tris, 150 mM NaCl and 0.1% Tween 20) and then incubated for at least 16 hours at 4°C with either mouse monoclonal anti-His (1:2,000; AD1.1.10, AbD Serotech) or mouse anti-RNA pol (1:5,000; 4RA2, Neoclone). Proteins were detected using anti-mouse IgG horseradish peroxidase (HRP)-conjugated secondary antibodies (1:3,000; catalog number NEF822001EA, Perkin-Elmer) and developed with Clarity Western ECL Substrate (BioRad). All antibodies were diluted in TBS-T with 1% bovine serum albumin (BSA; Sigma-Aldrich). Images were obtained using an MFChemiBis imaging station (DNR Bio-Imaging Systems) or an Amersham imager 600 (GE life sciences).

### Whole cell lysis of bacterial samples

Bacteria were grown overnight on LB plates. Plates were flooded with 5 ml of pre-chilled sterile phosphate-buffered saline (PBS) and colonies removed with a cell scraper. Cells were washed 3 times in PBS and collected by centrifugation at 10,000 × g at 4°C then snap frozen. Frozen whole cell samples were resuspended in 4% SDS, 100 mM Tris pH 8.0, 20 mM DTT and boiled at 95°C with shaking at 2000 rpm for 10 minutes. Samples were then clarified by centrifugation at 17,000 × g for 10 minutes, the supernatant collected, and protein concentration determined by bicinchoninic acid assay (Thermo Scientific Pierce). 200 μg of protein from each sample was acetone precipitated by mixing 4 volumes of ice-cold acetone with one volume of sample. Samples were precipitated overnight at -20°C and then centrifuged at 16,000 × g for 10 minutes at 0°C. The precipitated protein pellets were then resuspended in 80% ice-cold acetone and precipitated at -20°C. for an additional 4 hours. Samples were spun down at 17,000 × g for 10 minutes at 0°C to collect precipitated protein, the supernatant discarded, and excess acetone evaporated at 65°C for 5 minutes.

### Enrichment of DNA bound proteins

Bacteria were grown and washed as above then tumbled for 20 minutes at room temperature in 1% formaldehyde in PBS in accordance with the protocol of Qin *et al* (70). Cross-linking was quenched with 125 mM (final concentration) glycine in PBS for 5 minutes with tumbling. Cells were collected and washed twice with ice-cold PBS and then DNA isolated using a Zymo genomic DNA clean-up kit according to the manufacturer’s instructions with the exception of the DNA elution. DNA elution and reversal of DNA-protein cross-linking was undertaken by incubating the column at 68°C for 1 hour with protein elution buffer (2% SDS, 0.5 M β-mercaptoethanol and 300 mM Tris pH 8.5) according to Déjardin *et al* (71). Eluted proteins were collected by centrifugation and acetone precipitated by mixing 4 volumes of ice-cold acetone with one volume of sample. Samples were precipitated overnight at -20°C and then spun down at 16,000 × g for 10 minutes at 0°C. The precipitated protein pellets were precipitated with 80% ice-cold acetone and resuspended as indicated above.

### Peptidomic analysis of bacterial samples

Bacterial strains were grown and washed as above then solubilized by boiling in guanidinium chloride lysis buffer (6 M GdmCl, 100 mM Tris pH 8.5, 10 mM tris(2-carboxyethyl)phosphine, 40 mM 2-chloroacetamide) according to the protocol of Humphrey *et al* (72). Peptidomic samples were isolated according to Parker *et al* (73). Briefly, guanidinium chloride lysis buffer solubilised lysates were precipitated with 20% trichloroacetic acid on ice for 3 hours, then sample were centrifuged at 10,000 × g for 10 minutes at 0°C, the resulting supernatant collected and peptide isolated with tC18 columns (Waters). Bound peptides were eluted with 50% acetonitrile (ACN), 0.1% FA, dried and stored at -20°C.

### Digestion of complex protein lysates

Dried protein pellets were resuspended in 6 M urea, 2 M thiourea, 40 mM NH_4_HCO_3_ and reduced / alkylated prior to digestion with Lys-C (1/200 w/w) then trypsin (1/50 w/w) overnight as previously described (74). Digested samples were acidified to a final concentration of 0.5% formic acid and desalted with home-made high-capacity StageTips composed on 5 mg Empore™ C18 material (3M, Maplewood, Minnesota) and 5 mg of OLIGO R3 reverse phase resin (Thermo Fisher Scientific) according to the protocol of Ishihama and Rappsilber (75, 76). Bound peptides were eluted with Buffer B (80% ACN, 0.1% FA), dried and stored at -20°C.

### Reversed phase LC-MS

Purified peptides were resuspended in Buffer A* (2% ACN, 0.1% trifluoroacetic acid) and separated using a two-column chromatography set up comprising a PepMap100 C18 20 mm × 75 μm trap and a PepMap C18 500mm × 75μm analytical column (Thermo Scientific). Samples were concentrated onto the trap column at 5 μl/minute with Buffer A (2% ACN, 0.1%FA) for 5 minutes and infused into either an Orbitrap Elite™ Mass Spectrometer (Thermo Scientific), an Orbitrap Fusion Lumos Tribrid™ Mass Spectrometer (Thermo Scientific) or an Q-exactive plus™ Mass Spectrometer (Thermo Scientific) at 300 nl/minute via the analytical column using an Dionex Ultimate 3000 UPLC (Thermo Scientific). For whole cell proteomics analysis on the Orbitrap Elite™ 210 minute gradients were run altering the buffer composition from 1% buffer B to 28% B over 180 minutes, then from 28% B to 40% B over 10 minutes, then from 40% B to 100% B over 2 minutes, the composition was held at 100% B for 3 minutes, and then dropped to 3% B over 5 minutes and held at 3% B for another 10 minutes. The Orbitrap Elite™ was operated in a data-dependent mode automatically switching between the acquisition of a single Orbitrap MS scan (60,000 resolution) followed by 5 data-dependent HCD MS-MS events (resolution 15 k AGC target of 4 × 10^5^ with a maximum injection time of 250 ms, NCE 35) with 30 seconds dynamic exclusion enabled. For whole cell proteomics analysis on the Fusion™ 120 minute gradients were run altering the buffer composition from 1% buffer B to 28% B over 90 minutes, then from 28% B to 40% B over 10 minutes, then from 40% B to 100% B over 2 minutes, the composition was held at 100% B for 3 minutes, and then dropped to 3% B over 5 minutes and held at 3% B for another 10 minutes. The Fusion™ was operated in a data-dependent acquisition switching between the acquisition of an Orbitrap MS scan (120,000 resolution) every 3 seconds and HCD undertaken for each selected precursor (maximum fill time 100 ms, AGC 5 × 10^4^ with a resolution of 15,000 for Orbitrap MS-MS scans) with 30 seconds dynamic exclusion enabled.

DNA bound proteomic analysis was undertaken on both a Orbitrap Elite™ (for data-dependent acquisition experiments) and Q-exactive plus™ (for data-independent acquisition experiments) with 90 minute gradients run altering the buffer composition from 1% buffer B to 28% B over 60 minutes, then from 28% B to 40% B over 10 minutes, then from 40% B to 100% B over 2 minutes, the composition was held at 100% B for 3 minutes, and then dropped to 3% B over 5 minutes and held at 3% B for another 10 minutes. For data-dependent acquisition experiments the Elite™ Mass Spectrometer was operated in a data-dependent mode automatically switching between the acquisition of a single Orbitrap MS scan (60,000 resolution) followed by 10 data-dependent CID MS-MS events (analysed within the ITMS, maximum injection time of 100 ms, NCE 35). To enable the robust quantification of CepR data-independent acquisition was undertaken using parallel reaction monitoring (PRM (77)) monitoring tryptic peptides of BCAL3530 (DNA-binding protein HU-alpha), BCAL0462 (Putative DNA topoisomerase III), BCAM0904 (DNA polymerase I), BCAM1868 (CepR) and BCAM1870 (CepI), Supplementary Table 4. BCAL3530, BCAL0462, BCAM0904 were included as positive controls to ensure equal DNA enrichment while CepI is a non-DNA binding control. Data-independent acquisition was performed by switching between the acquisition of a single Orbitrap MS scan (70,000 resolution, m/z 350-1400) and HCD MS/MS events of each PRM precursor (maximum fill time 110ms, AGC 2 × 10^5^ with a resolution of 35,000 for Orbitrap MS-MS scans, see Supplementary Table 4 for further details).

Peptidomic analysis was undertaken on an Orbitrap Elite™ with 120 minute gradients run altering the buffer composition from 1% buffer B to 28% B over 90 minutes, then from 28% B to 40% B over 10 minutes, then from 40% B to 100% B over 2 minutes, the composition was held at 100% B for 3 minutes, and then dropped to 3% B over 5 minutes and held at 3% B for another 10 minutes. The Elite™ Mass Spectrometer was operated in a data-dependent mode automatically switching between the acquisition of a single Orbitrap MS scan (60,000 resolution) followed by 5 data-dependent CID and HCD MS-MS events (both acquired within the Orbitrap for each precursor at a resolution of 15 k AGC target of 4 × 10^5^ with a maximum injection time of 250 ms, NCE 35) with 30 seconds dynamic exclusion enabled.

### Data analysis

MS datasets were processed using MaxQuant (v1.5.5.1 or 1.5.3.30 (78)). Database searching was carried out against the reference *B. cenocepacia* strain J2315 (https://www.uniprot.org/proteomes/UP000001035, downloaded September 25^th^ 2017) and the K56-2Valvano proteome (79) (http://www.uniprot.org/taxonomy/985076, downloaded from NCBI February 15^th^ 2013). All tryptic digest searchers were undertaken using “Trypsin” enzyme specificity, carbamidomethylation of cysteine as a fixed modification; oxidation of methionine, acetylation of protein N-terminal trypsin/P cleavage with a maximum of 2 missed cleavages. Peptidomic analysis was undertaken using “Unspecific” enzyme specificity and allowing the presence of *O*-linked glycosylation BC glycan 1 (elemental composition: C_22_O_15_H_36_N_2_, mass: 568.2115) and BC glycan 2 (elemental composition: C_26_O_18_H_40_N_2_, mass: 668.2276) at serine residues. To enhance the identification of peptides between samples, the Match between Runs option was enabled with a precursor match window set to 2 minutes and an alignment window of 10 minutes. For label free quantitation the MaxLFQ option in Maxquant (80) was enabled in addition to the re-quantification module. The resulting outputs were processed within the Perseus (v1.5.0.9) (81) analysis environment to remove reverse matches and common protein contaminates prior to further analysis. For label-free based quantitative (LFQ) comparisons missing values were imputed using Perseus and data z-scored to enabled visualization in heat maps using R (https://www.r-project.org/). Pearson correlations were performed on non-imputed dataset with clustering and visualization performed in Perseus. Clustering of Pearson correlations data was undertaken using the Euclidean distance and complete linkage with pre-processing with k-means in Perseus. Enrichment analysis was undertaken using Fisher exact test in Perseus with Gene Ontology (GO) terms, gene names, subcellular location [CC], signal peptide status, lipidation status, intramembrane status and keywords associated with each protein obtain from uniport (*B. cenocepacia* strain J2315 proteomes: UP000001035, downloaded September 25^th^ 2017). Virulence associated genes were compiled from genes defined as Virulence associated within the *Burkholderia* Genome Database (82) (J2315, Downloaded 3*^rd^* October 2017) and the genome analysis of *B. cenocepacia* strain J2315 (83) (Supplementary Table 6). CepR regulated proteins were defined as those proteins which were previously reported by O’Grady *et al* as differential regulated in K56-2 Δ*cepR* at stationary phase (43). Proteomics data sets have been deposited to the ProteomeXchange Consortium via the PRIDE (84) partner repository with the dataset identifier PXD014429, PXD014516, PXD014581, PXD014614 and PXD014700. For a complete description of each PRIDE dataset see Supplementary Table 5.

### Motility assays

Motility assays were conducted using semi-solid motility agar consisting of LB infusion medium supplemented with 0.3 % agar as previously described (56). Plates were inoculated using 2 μl of standardized, OD_600_ of 0.5, overnight cultures of each strain. Motility zones were measured after 48 hour incubation at 37°C. Experiments were carried out in triplicate with 3 biological replicates of each strain.

### Transcriptional Analysis by luminescence assays

To assess transcriptional changes in CepR and CepI *luxCDABE* reporter assays were performed using *B. cenocepacia* K56-2 wild-type (WT), Δ*pglL*, Δ*OGC* and Δ*pglL amrAB::S7-pglL-his*_10_ strains containing pCP300 (CepI promoter *luxCDABE* reporter (85)), pPromcepR (CepR promoter *luxCDABE* reporter (86)) or pMS402 (promoterless *luxCDABE* reporter (87)) as a negative control.

Overnight cultures were diluted to an OD_600_ of 1.0 and 2 μl inoculated into 200 μl LB supplemented with 100 μg/ml trimethoprim in black, clear-bottomed 96-well microplates (minimum of eight technical replicates per independent biological replicate). The OD_600_ and relative luminescence were measured using a CLARIOstar plate reader at 10-minute intervals for 24 hours. Experiments assessing the effect of C_8_-HSL additions on CepR and CepI transcription were performed according to Le Guillouzer *et al* (88). Briefly, cultures were supplemented with C_0_-HSL (Sigma-Aldrich) resuspended in acetonitrile (10 µM final concentration) and added to cultures with acetonitrile added alone used as a negative control. Plates were incubated at 37°C with shaking at 200 rpm between measurements with each assay undertaken 3 independent times on separate days. The resulting outputs were visualised using R (https://www.r-project.org/).

### Biofilm Assay

Biofilm assays were performed according to previous reports (26, 89, 90) using protocols based on the approach of O’Toole (91). *B. cenocepacia* strains were grown overnight at 37°C and adjusted to an OD600 of 1.0. 10 ml of these suspensions were inoculated into 990 ml of LB supplemented with 0.5% (wt/vol) casamino acids and 100 ml added in to 96-well microtiter plates (Corning Life Sciences, minimum of eight technical replicates per independent biological replicate). Microtiter plates were incubated at 37°C for 24 hours in a closed humidified plastic container. The plates were then washed with PBS to remove planktonic cells then stained for 15 minutes with 125 μl of 1% (wt/vol) crystal violet. Excess crystal violet was removed with two washes of PBS and 200 μl of 33% (vol/vol) acetic acid was added for 15 minutes to release the stain. Resuspended stain was transferred to a new plate and measured on a CLARIOstar plate reader measuring the absorbance of the resulting solution at 595 nm. Three independent assays were undertaken on separate days.

### Galleria mellonella infection assays

Infection of *G. mellonella* larvae was undertaken using the approach of Seed and Dennis (92) with minor modifications. *B. cenocepacia* strains were grown overnight at 37°C and adjusted to an OD_600_ of 1.0, equivalent to 2 × 10^9^ cfu/ml (colony-forming units / ml). Strains were diluted with PBS to 4 × 10^5^ cfu/ml with serial dilution plates undertaken to confirm inoculum levels. For each strain, 2000 cfu in 5 μl was injected in the right proleg of the *G. mellonella* larvae. 3 independent challenges were performed with each strain injected into 8 to 10 *G. mellonella* larvae. For each independent challenge 8 control larvae were injected with 5 μl PBS. Post infection *G. mellonella* larvae were placed in 12-well tissue culture plates and incubated in the dark at 30°C. The number of dead larvae was scored at 24, 48, and 72 hours after infection with death of the larvae determined by loss of responsiveness to touch. The results visualised using R (https://www.r-project.org/) and statistical analysis of survival curves undertaken with the survminer package (version 0.4.5).

### CAS siderophores assays

Alterations in production of siderophores was assessed using Chrome Azurol S (CAS) assay as previously described (93, 94). 10 μl of OD_600_ 1.0 adjusted bacterial culture were spotted on CAS agar plates and incubated at 37°C for 24 hours. The diameter of the zone of discoloration from the removal of iron from the CAS dye complex was measured. Experiments were carried out in technical triplicate with 3 biological replicates.

### Proteases activity-based probes

K56-2 WT, Δ*pglL* and Δ*pglL amrAB::S7-pglL-his*_10_ strains were grown overnight on confluent LB plates. Plates were flooded with 5 ml of pre-chilled sterile PBS and colonies removed with a cell scraper. Cells were washed 3 times in chilled PBS and resuspend in 40 mM Tris, 150 mM NaCl pH 7.8 then lysed by sonication. Samples were clarified by centrifugation, 10,000 × g, 10 minutes at 4°C and samples were diluted to a total concentration of 4 mg/ml. Reactivity to three classes of activity-based probes were assessed using PK-DPP (Cy5-tagged probe for Trypsin-like proteases (95)), PK105b (Cy5-tagged probe for Elastase-like proteases (96)), PK101 (Biotin-tagged probe for Elastase-like proteases (97)) and FP-Biotin (Biotin-tagged probe for Serine hydrolases (98)) which were added at 1.3 µM from a 100x DMSO stock. Untreated control samples were prepared in parallel and left untreated to allow the assessment of autofluorescence and endogenous biotinylation in lysates. Samples were incubated at 37°C for 15 minutes to allow labelling and then quenched by the addition of Laemmli Sample Buffer. Samples were then boiled and proteins were resolved on a 15% SDS-PAGE gel. For Cy5-tagged probes, labelling was detected by directly scanning the gel for Cy5 fluorescence using a Typhoon 5 flatbed laser scanner (GE Healthcare). For FP-Biotin, proteins were transferred to nitrocellulose and the membrane was incubated with streptavidin-AlexaFluor647 at 4°C overnight. Following three washes with PBS containing 0.05% Tween 20, the membrane was scanned on the Typhoon 5 in the Cy5 channel. Experiments were carried out in biological triplicate. All probes were synthesised in house by the Edgington-Mitchell Laboratory according to published methods, with the exception of FP-Biotin, which was purchased from Santa Cruz Biotechnology.

## Results

### Loss of glycosylation in *B. cenocepacia* leads to global proteome alterations

We previously demonstrated that loss of glycosylation causes defects in motility (56), reduction of virulence in plant and insect infection models (56, 65), and defects in carbon utilisation (65). To better understand the role of glycosylation in *B. cenocepacia* we assessed the effect of loss of glycosylation on the proteome. To achieve this we generated markerless deletion mutants in the *O*-oligosaccharyltransferase *pglL (ΔpglL*; BCAL0960 (56)) gene, the recently identified *O*-linked glycan cluster *(ΔOGC*; BCAL3114 to BCAL3118) responsible for the generation of the glycan used for *O*-linked glycosylation (65), and a double glycosylation null strain *(ΔpglLΔOGC*). We also constructed a chromosomal *pglL* complemented strain *(ΔpglL amrAB::S7-pglL-his*_10_, Supplementary Figure 1A). The rationale for creating multiple glycosylation-defective strains was to eliminate potential confounding effects arising from blocking glycosylation at a specific step and the corresponding accumulation of unprocessed lipid-linked glycans. Western blot analysis using the glycoprotein acceptor protein DsbA_Nm_-his_6_ (56, 99) confirmed the loss of glycosylation in Δ*pglL*, Δ*OGC*, and Δ*pglLΔOGC*, as well as restoration of glycosylation in Δ*pglL amrAB::S7-pglL-his*_10_ (Figure 1A). In contrast to our previously reported plasmid based PglL complementation approaches (56) chromosomal complementation lead to the restoration of glycosylation to near wild-type levels (Supplementary Figure 1B) as well as restoration of motility (Supplementary Figure 1C) compared to only partial restoration previously reported (56).

**Figure 1.**
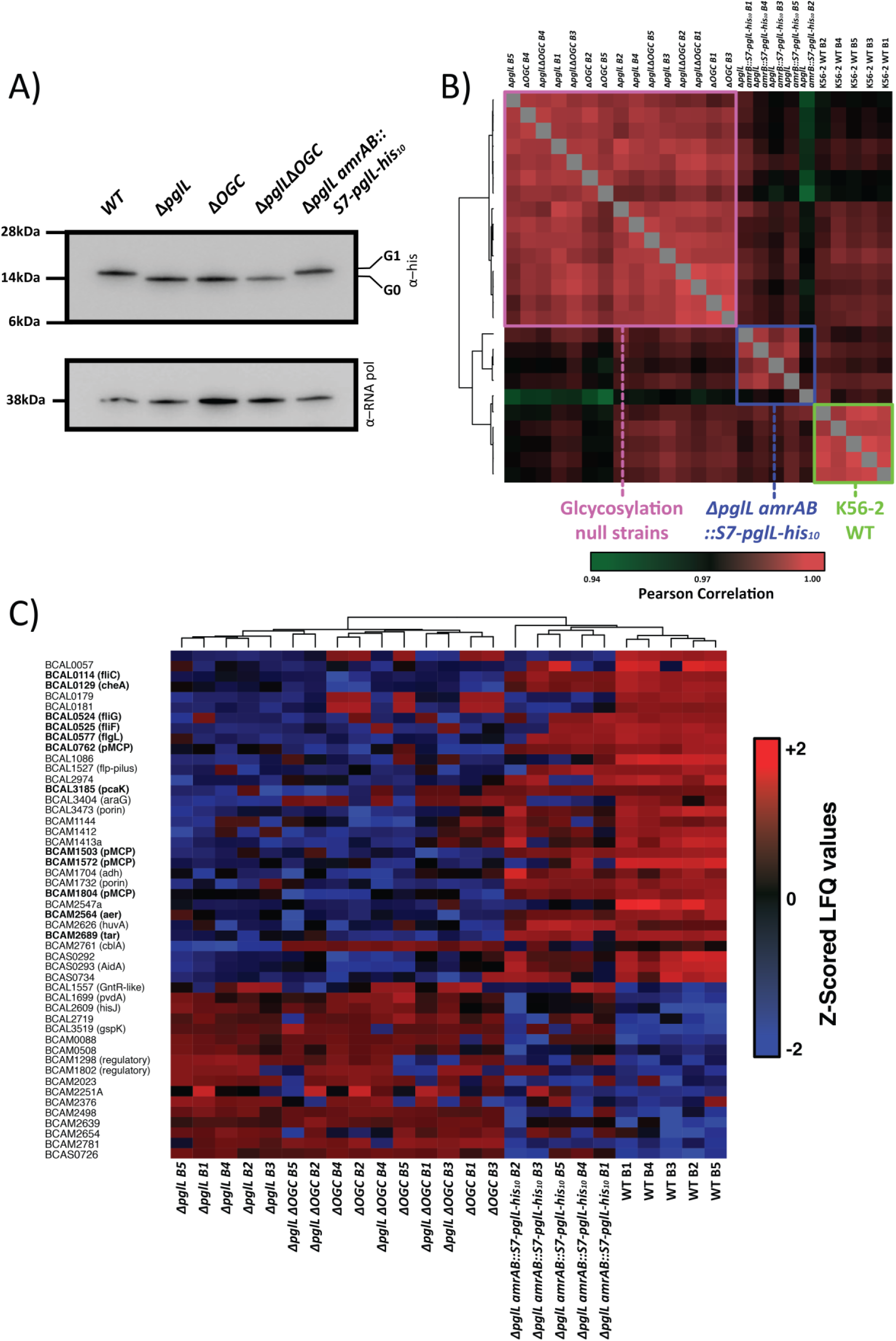
Disruption of *O*-linked glycosylation results in multiple changes in the proteome. A) Western analysis of strains expressing the glycosylation substrate DsbA_Nm_-his_6_ confirms the loss of glycosylation in Δ*pglL*, Δ*OGC*, Δ*pglLΔOGC* and restoration of glycosylation in the chromosomal complement Δ*pglL amrAB::S7-pglL-his*_10_. B) Pearson correlation analysis demonstrate three discrete clusters observed across the proteomic analysis which separate glycosylation competent and glycosylation-null strains. C) Z-scored heatmap of proteins observed to undergo alterations between glycosylation-competent and glycosylation-null strains reveals alterations in motility and chemotaxis (proteins bolded) including BCAL0114 (FliC), BCAL0524 (FliG), BCAL0525 (FliF) as well as known CepR regulated protein BCAS0293 (AidA).

Using label-free based quantitative proteomics, 5 biological replicates of each strain were investigated leading to the identification of 3399 proteins with 2759 proteins quantified in at least 3 biological replicates in a single biological group (Supplementary Figure 2, Supplementary Table 7). Hierarchical clustering of Pearson correlations of proteome samples demonstrated robust correlation between all samples (Average Pearson correlation of 0.98, Supplementary table 8) yet three discrete proteome clusters were readily identified separating the wild-type K56-2, Δ*pglL amrAB::S7-pglL-his*_10_ strains and glycosylation-null strains (Figure 1B). Examination of the most profound alterations, proteins with –log_10_(p)>3 and a fold change greater than +/-2 log_2_ units, revealed alterations in protein levels observed in Δ*pglL* that were mirrored in strains Δ*OGC* and Δ*pglLΔOGC*, which were restored by complementation (Figure 1C). Consistent with the observed motility defects (Supplementary Figure 1C), the levels of proteins associated with flagella-mediated motility and chemotaxis, including BCAL0114 (FliC), BCAL0129 (CheA), BCAL0524 (FliG) and BCAL0525 (FliF), were significantly reduced in glycosylation-null strains. Importantly, multiple known virulence-associated proteins were also decreased in the glycosylation-null strains including the heme receptor protein BCAM2626 (HuvA (100)) and nematocidal protein BCAS0293 (AidA (101)). Numeration of the overlap of all altered protein between glycosylation-null strains by Fisher exact enrichment analysis demonstrated a substantial enrichment between these three groups (Fisher exact test: 6.7502 × 10^177^ and 4.3784 × 10^245^ for Δ*pglL* compared with Δ*OGC*, and for Δ*pglL* compared with Δ*pglLΔOGC*, respectively, Supplementary Table 9, Supplementary Figure 3). These results revealed that the loss of glycosylation due to disruption of *pglL* or *OGC* leads to similar changes, which are largely complemented to parental levels by reintroduction of *pglL* in the chromosome.

### Loss of glycosylation results in reduced CepR/I transcription and the levels of DNA associated CepR

Enrichment analysis of the altered proteins within glycosylation-null strains demonstrate the over representation of a range of categorical groups based on GO terms, protein localization and virulence associated factors assignments. These groups highlight that protein localizations assignments and virulence associated factors were similarly affected in Δ*pglL* and Δ*OGC*, recapitulating observations made at the individual protein level (Figure 2, Supplementary Table 9). Interestingly, enrichment analysis highlighted the link between the loss of *O*-linked glycosylation and changes that were broader than only motility and virulence. For example, differences also observed in proteins associated with DNA-sequence specific binding and transcriptional regulation (Figure 2, Supplementary Table 9). This observation suggested that the loss of glycosylation results in alterations in the transcriptional landscape of *B. cenocepacia*. As virulence is coordinated by global regulators such as CciR, CepR, ShvR and AtsR in *B. cenocepacia* (35, 43, 86, 102), we assessed if known regulators could account for the observed proteome changes in glycosylation-null strains. As our data demonstrated minimal alteration of the regulator ShvR (BCAS0225, Supplementary Table 7) across the analysed strains, and disruption of both *atsR* (*BCAM0379*) and *cciR* (*BCAM0240*) has previously been associated with increased motility (43, 86), we reasoned that the regulator CepR may be responsible for the glycosylation-dependent differences in our mutant strains. This would be consistent with the observed decrease in BCAS0293 (AidA, Figure 1C) which is stringently regulated by CepR (45, 103). Using available microarray data of CepR-regulated genes (43), we investigated the correlation of the proteome changes observed in the absence of glycosylation, with alterations observed in response to the disruption of CepR. We observed a statistically significant enrichment of CepR-regulated proteins altered in the absence of glycosylation (multiple hypothesis corrected *p*-value 1.79 × 10^6^ and 6.69 × 10^6^ for Δ*pglL* and Δ*OGC* respectively, Supplementary table 9), supporting a link between CepR and the alteration observed in glycosylation-null strains and suggesting that the loss of glycosylation may influence the *B. cenocepacia* CepR regulon.

**Figure 2.**
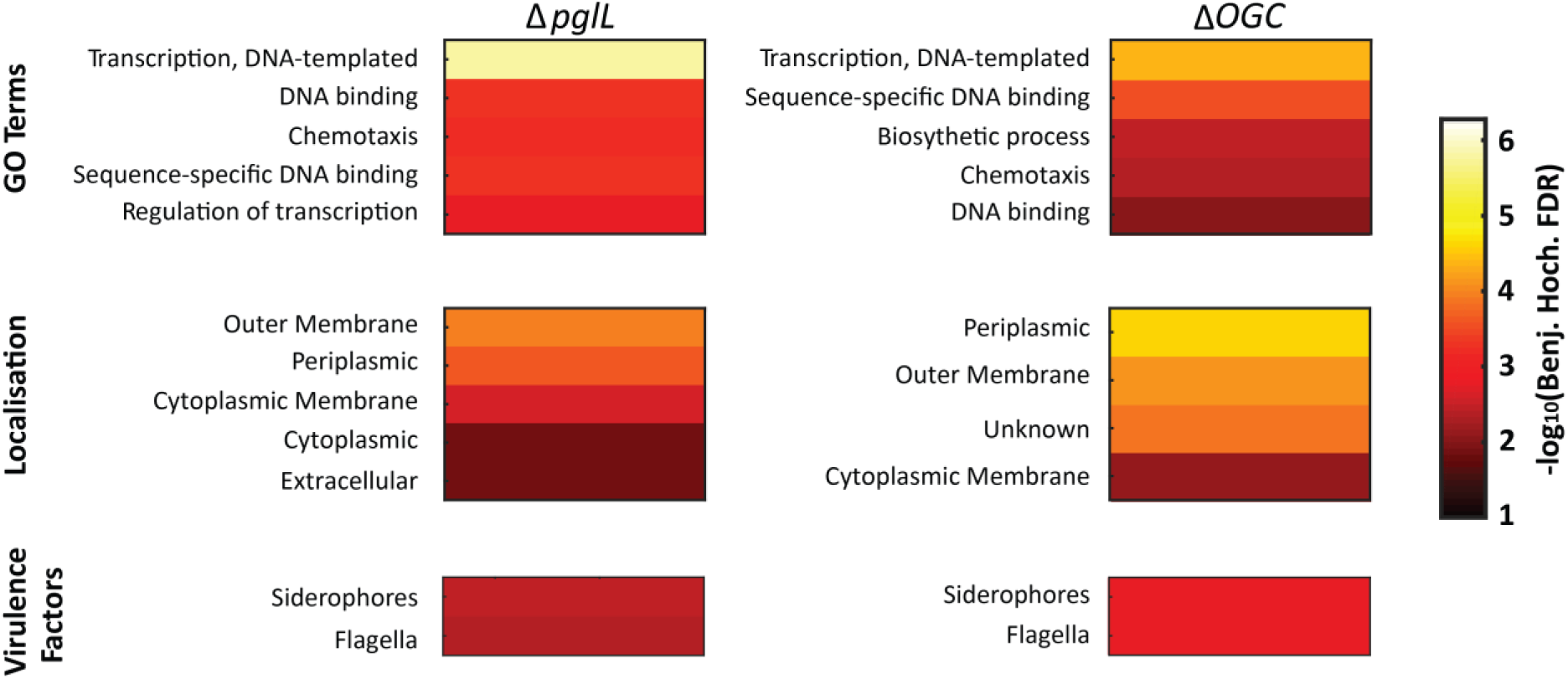
Heatmaps of glycosylation-null strains enrichment analysis. The multiple hypothesis corrected *p*-values from Fisher exact tests demonstrate proteins with similar GO terms, localisations and associated with virulence factors are altered with glycosylation-null strains.

To determine transcriptional changes in *cepR/I* genes we introduced the *cepR* and *cepI* luciferase promoter reporter (pPromcepR (86) and pCP300 (85)) into wild-type K56-2, Δ*pglL* and Δ*pglL amrAB::S7-pglL-his*_10_. Consistent with our proteomic results, Δ*pglL* showed decreased induction of both *cepI* and *cepR* over a 24-hour period (Figure 3A, Supplementary Figure 4) compared with wild type and Δ*pglL amrAB::S7-pglL-his*_10_. Detailed examination at 12 hours (log phase), 16 hours (the transition from log to stationary phase) and 20 hours (stationary phase) revealed higher levels of transcription in the wild type of both *cepI* and *cepR* at 16 and 20 hours compared with transcription levels in Δ*pglL*, despite comparable growth kinetics (Supplementary Figure 5). As the C8-HSL levels affect the response of CepI and CepR in *B. cenocepacia* (39, 44, 104), we assayed *cepR/I* transcription in the absence and presence of additional C8-HSL (10 μM, Figure 3B). In response to exogenous C8-HSL, *cepI* transcription increased in all strains (Figure 3B), consistent with the positive feedback response expected to heighten C8-HSL levels (39, 44). In contrast, while the addition of C8-HSL led to no change in *cepR* transcription in Δ*pglL*, it resulted in reduced transcription of *cepR* to the level observed in the wild type K56-2. As expected from the reduction in *cepR/I* transcription resulting from the loss of glycosylation, *cepR* and *cepI* transcription was also compromised in Δ*OGC* strains (Supplementary Figure 6 and 7). Together, these results indicate that both *cepR* and *cepI* transcription are altered in the loss of glycosylation, with the resulting *cepR* levels resembling the levels observed during C8-HSL-induced repression in wild type.

**Figure 3.**
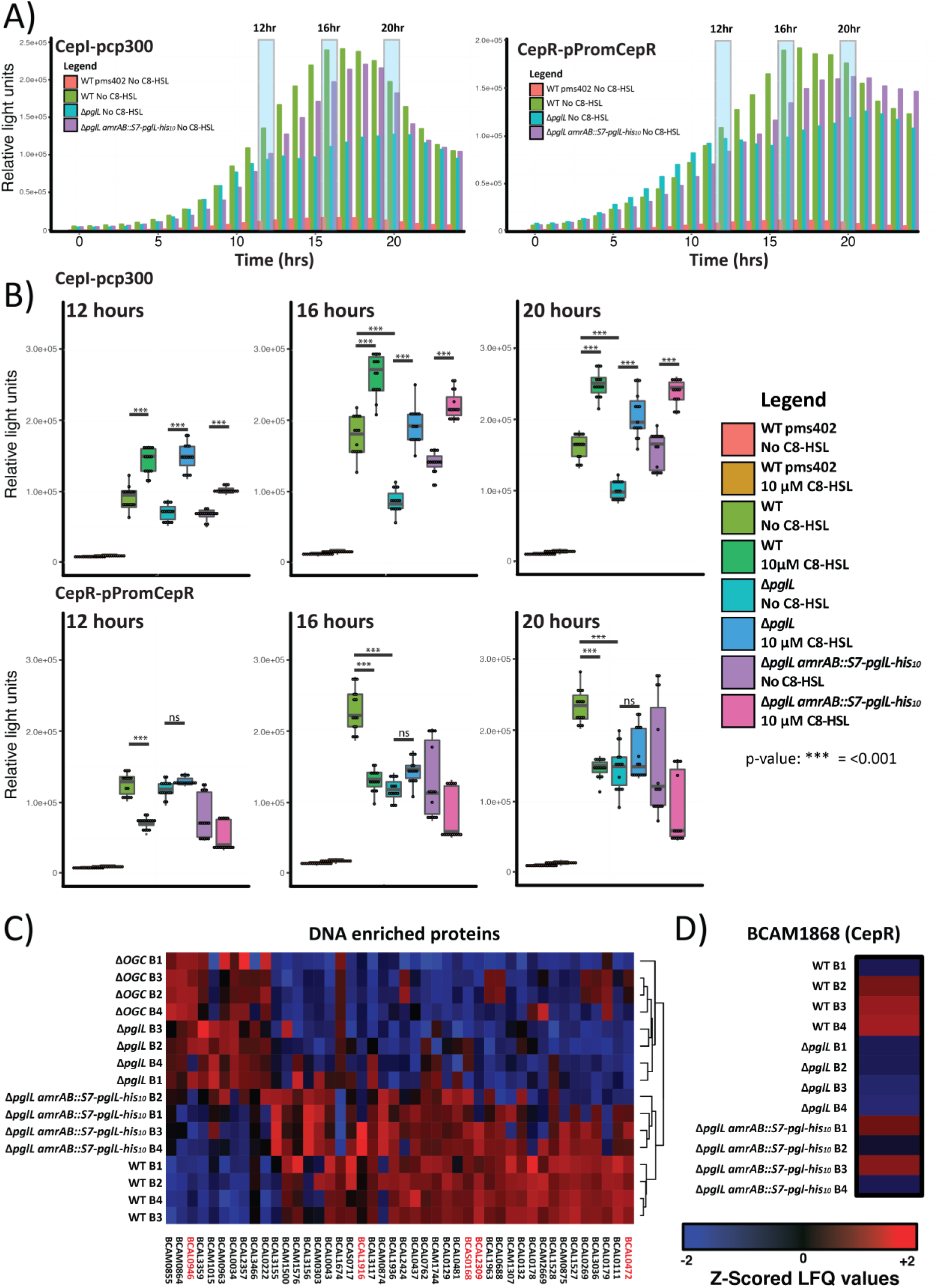
CepR/I transcription are altered in glycosylation-null strains. A) 24-hour luciferase profile of strains grown with either the CepI reporter pCP300 or CepR reporter pPromCepR demonstrating alteration in luciferase activity in the Δ*pglL* compared to WT and Δ*pglL amrAB::S7-pglL-his*_10_ strains. Each data point corresponds to the mean of three independent biological replicates with a more detailed figure containing the plotted standard deviation provided in Supplementary Figure 4. B) Detailed analysis of three time points across the luciferase profiles are provided for the 12-hour (log), 16-hour (transition from log to stationary) and 20-hour (stationary) time points. For each time point the luciferase activity with strains grown with and without the presence of C8-HSL are shown. C) Z-scored heatmap of DNA bound proteins with significant alterations in abundance in Δ*pglL* or Δ*OGC* compared to WT reveal similar protein profiles for glycosylation-null strains compared to glycosylation-competent strains. D) DNA bound proteome analysis of CepR supports the reduction in the abundance of DNA bound CepR in Δ*pglL* and the partial restoration of CepR in Δ*pglL amrAB::S7-pglL-his*_10_.

As the CepR protein autoregulates *cepR*’s own transcription (48), we reasoned that the decreased transcription in Δ*pglL* would correspond to decreased levels of DNA bound CepR. To directly assay DNA binding by CepR, we monitored the DNA bound proteome using formaldehyde-based cross-linking coupled to DNA enrichment (105). Initial analysis of the DNA-bound proteome found glycosylation-null strains (Δ*pglL* and Δ*OGC*) and glycosylation-proficient strains (wild type and Δ*pglL armAB::S7-pglL-his*_10_) possessed distinct proteome profiles with multiple uncharacterised transcriptional regulators (e.g. BCAL0946, BCAL1916, BCAS0168, BCAL2309 and BCAL0472) which were altered by the loss of glycosylation (Figure 3C, Supplementary Table 10). Although this analysis enabled the identification of CepR, its low abundance prevented its quantitation across biological replicates. To improve the monitoring of CepR, targeted proteomic analysis was undertakening using PRM assays confirming the reduction in DNA associated CepR in Δ*pglL* compared with wild type and Δ*pglL armAB::S7-pglL-his*_10_ (Figure 3D, *p*-value= 0.017 wild type vs ΔpglL, Supplementary Table 11). In agreement with the total proteome and *lux* reporter measurements, the DNA-bound proteome supports multiple transcription associated proteins, including the global regulator CepR, that are altered in the absence of glycosylation.

### Δ*pglL* demonstrates a reduced ability to form biofilms and produce siderophores

The observed reductions in CepR/I transcription suggested that CepR/I-linked regulation may also be altered in glycosylation-null strains. To test this hypothesis, we assessed two phenotypes associated with CepR/I regulation, the production of biofilm under static 24-hour growth and siderophore production (39, 43–45, 48). Consistent with an impact of glycosylation on CepR/I-mediated regulation, we observed a profound reduction in biofilm formation in Δ*pglL*, which was partially restored by complementation (Figure 4A). Interestingly, we observed that the method of complementation, i.e. expression of PglL-his_10_ driven from the native pglL promoter (Δ*pglL amrAB::native-pglL-his*_10_) or from the constitutive S7 promoter (Δ*pglL amrAB::S7-pglL-his*_10_) impacted the restoration of biofilm formation (Figure 4A). Examination of independently created Δ*pglL* and Δ*pglL amrAB::native-pglL-his*_10_ strains confirmed a link between biofilm formation, through phenotype restoration following complementation (Supplementary Figure 8). CAS assays, used to assess the global levels of siderophore activity, demonstrated a modest but reproducible effect in Δ*pglL* which was completely restored by complementation when PglL was expressed from either the native or S7 promoter (Figure 4B and C). Together, these findings support the notion that CepR/I regulate multiple phenotypes including biofilm and siderophore production, which are affected by the loss of glycosylation.

**Figure 4.**
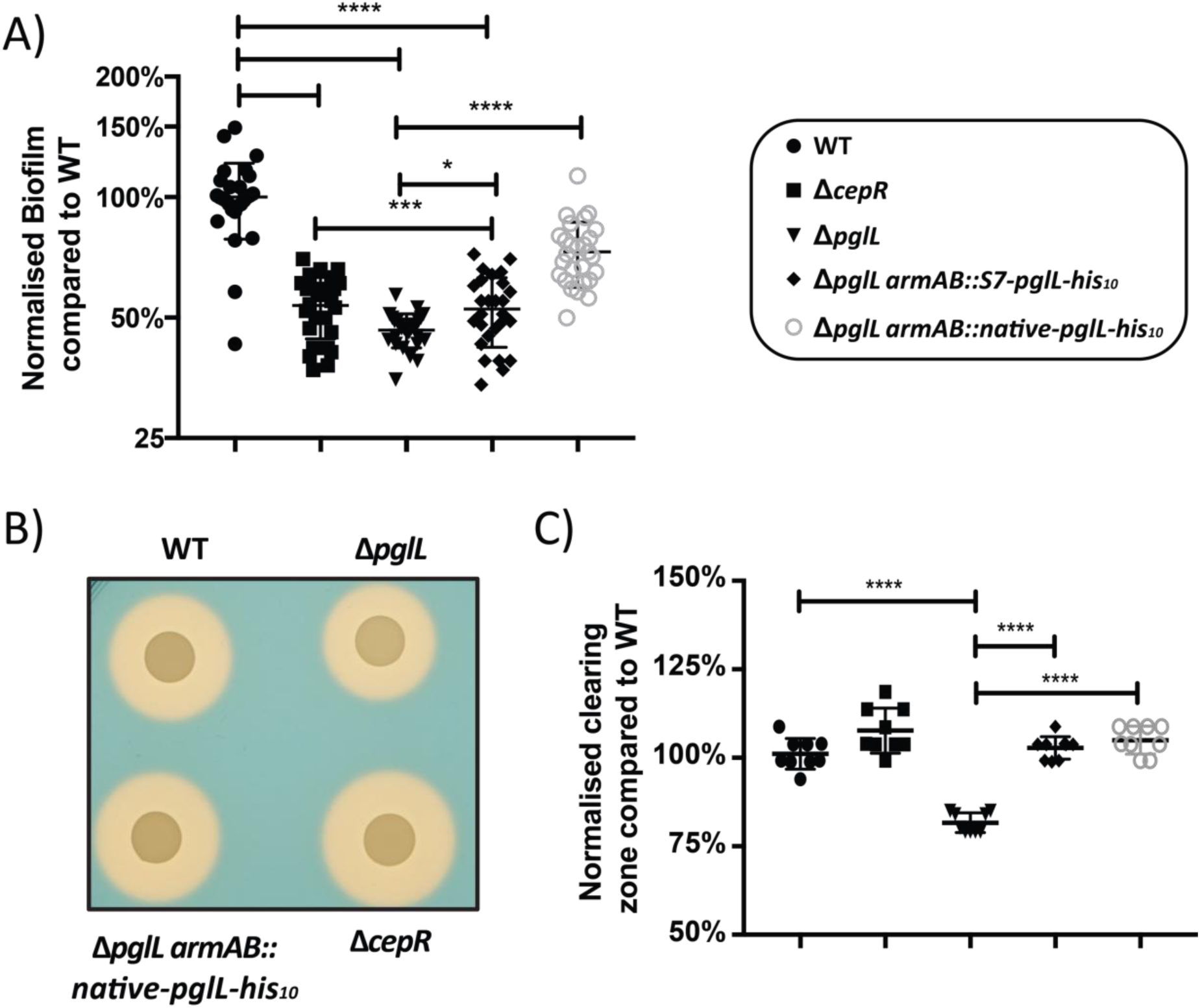
Biofilm formation and siderophore production are reduced in Δ*pglL*. A) 24-hour static biofilm assays demonstrate a decrease in biofilm formation in Δ*pglL* which is partially restored upon complementation. B and C) CAS assays demonstrates a reduction in zone of clearing in Δ*pglL* which is restored upon complementation.

### Except for BCAL1086 and BCAL2974, proteins that are normally glycosylated remain stable in the absence of glycosylation

As the loss of glycosylation in other bacterial glycosylation systems leads to protein instability (63, 64, 106), we examined whether protein instability in *B. cenocepacia* may be responsible for the phenotypic changes in glycosylation-null strains. Our proteomic analysis identified 21 out of 23 known glycoproteins (56), yet only 2 were altered in abundance in glycosylation-negative strains; BCAL1086 (-5.7 log_2_) and BCAL2974 (-2.5 log_2_) (Figure 5A, Supplementary Table 7). To confirm the observed decreases in abundance, endogenous BCAL1086 and BCAL2974 were his_10_-tagged at the C’-terminus. While his-tagging did not allow the detection of BCAL2974 by western analysis (data not shown), the introduction of the his_10_ epitope into BCAL1086 allowed quantification of endogenous BCAL1086 in the K56-2 wild-type, Δ*pglL* and Δ*pglL amrAB::S7-pglL-his*_10_ backgrounds and confirmed the loss of BCAL1086 in Δ*pglL* (Figure 5B). We sought to directly assess whether BCAL1086 was subjected to increased degradation in Δ*pglL*, as a measure of instability. For this, we monitored the endogenous peptide pool (73) quantifying peptides derived from 783 proteins (Supplementary Table 12 and 13) in *B. cenocepacia* K56-2 WT, Δ*pglL* and Δ*pglL amrAB::S7-pglL-his*_10_ strains. Consistent with the degradation of BCAL1086 we observed an increase in the abundance of BCAL1086-derived peptides in Δ*pglL* while peptides from other known glycoproteins showed only modest changes (Figure 5C, Supplementary Table 13). Within this peptidomic analysis, we observed that multiple unique BCAL1086 peptides are were present in Δ*pglL* clustered around the central region of BCAL1086 (Figure 5D), confirming that BCAL1086 was expressed in Δ*pglL* but was subject to proteolysis. Together, our data support that BCAL1086 becomes degraded in the absence of glycosylation, but the majority of known *B. cenocepacia* glycoproteins are unaffected.

**Figure 5.**
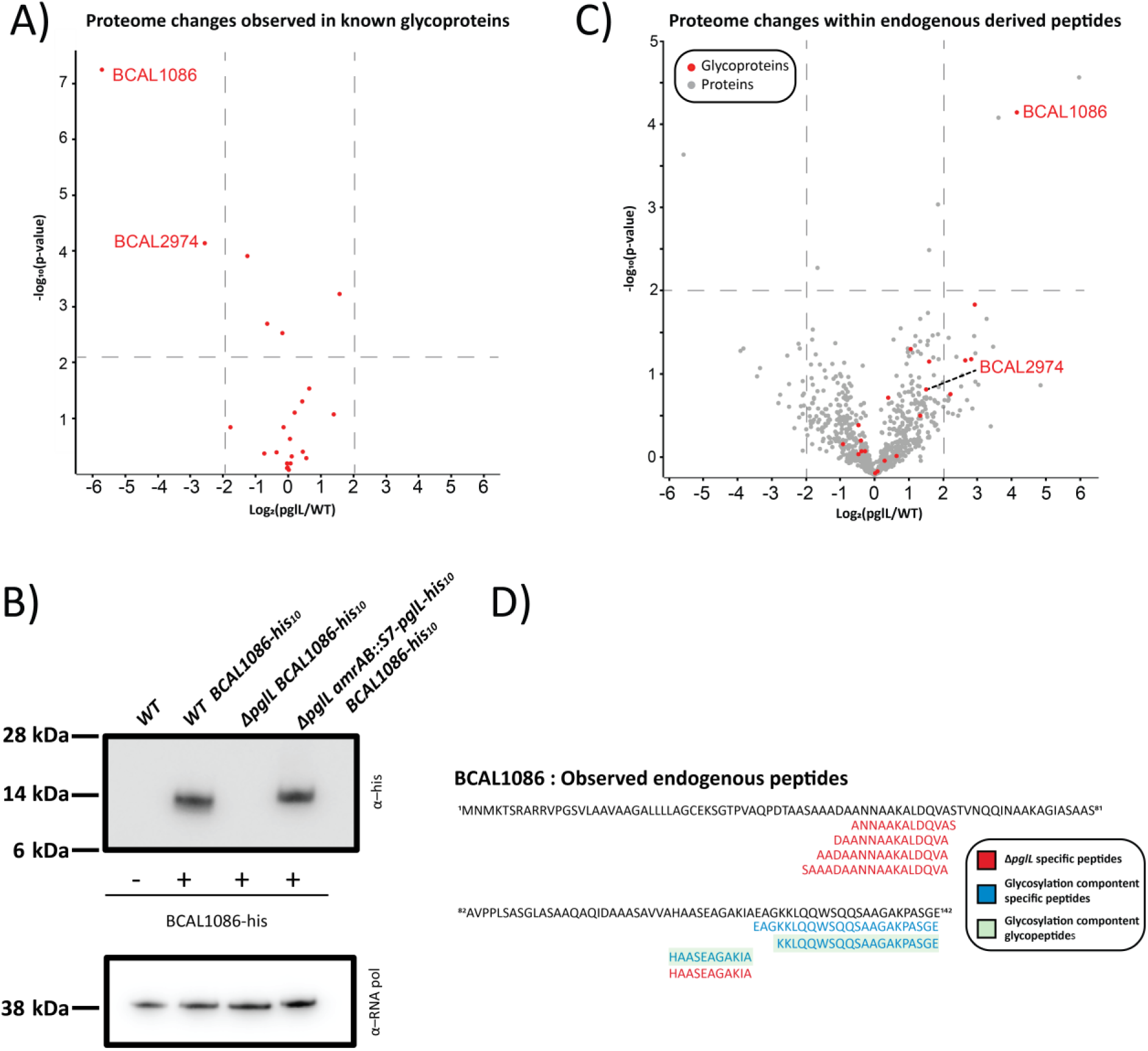
The stability of the glycoprotein BCAL1086 and BCAL2974 are affected by loss of glycosylation. A) Proteomic analysis demonstrates BCAL1086 and BCAL2974 decrease in abundance in the absence of glycosylation. B) Endogenous tagging of BCAL1086 confirms the loss of BCAL1086 in the Δ*pglL* background. C) Proteomics analysis of endogenous derived peptides demonstrates an increased abundance of BCAL1086-derived peptides in the absence of glycosylation. D) Analysis of endogenous peptides confirms the presence of unique peptide fragments from BCAL1086 in the Δ*pglL* background.

### Role of BCAL1086 and BCAL2974 in Δ*pglL* phenotypes

As changes in the glycoproteins BCAL2974 and BCAL1086 coincided with an alteration in CepR/I activity, we investigated if the loss of BCAL2974 and BCAL1086 could be responsible for defects observed in Δ*pglL*. To answer this question, Δ*BCAL2974* and Δ*BCAL1086* strains were created and assessed for their effect on biofilm and siderophore production as well as virulence in *G. mellonella*, a phenotype previously associated with Δ*pglL* (56). Assessment of 24-hour static biofilm growth showed Δ*BCAL1086* had no effect on biofilm formation while Δ*BCAL2974* resulted in a small but consistent decrease in biofilm development. However, this effect is minimal compared to the defect observed in Δ*pglL* and Δ*cepI* (Figure 6A). The ability of Δ*BCAL1086* and Δ*BCAL2974* to produce siderophores was unaffected (Figure 6B and C). Similarly, while *G. mellonella* infections showed that Δ*pglL* causes reduced mortality at 48 hours post infection compared to K56-2 WT, (*p*-value 0.0015), Δ*BCAL2974*, Δ*BCAL1086* and Δ*pglL amrAB::native-pglL-his*_10_ strains resulted in wild type levels of lethality in *G. mellonella* at 48 hours (Figure 6D). These results suggest that even though BCAL2974 and BCAL1086 are influenced by the loss of glycosylation, neither protein is solely responsible for the known defect observed in ΔpglL.

**Figure 6.**
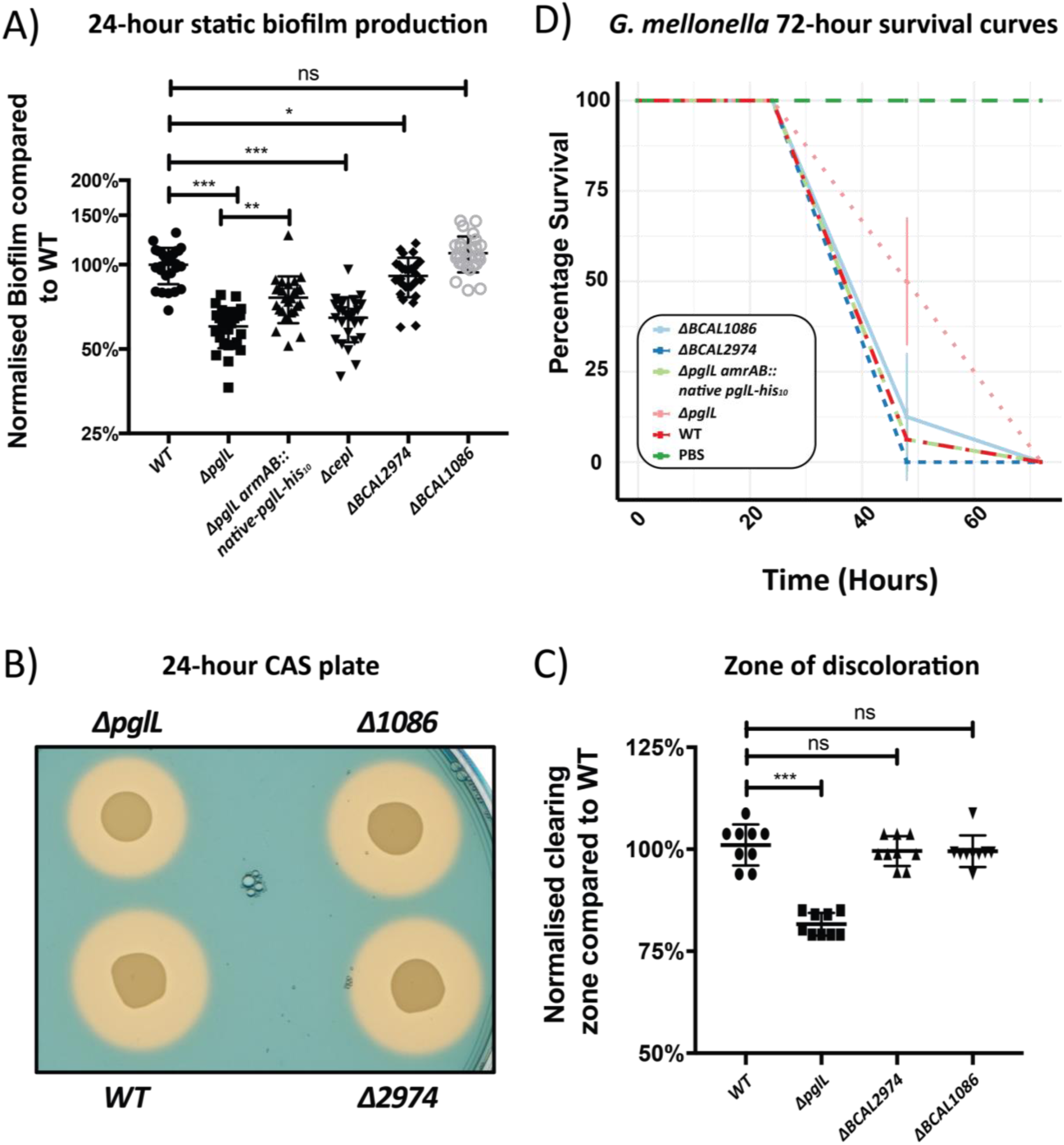
The loss of BCAL1086 or BCAL2974 does not affect phenotypes associated with ΔpglL. A) 24-hour static biofilm formation is unaffected in Δ*BCAL1086* and minimally affected in Δ*BCAL2974* compared to WT. B) CAS plate assays demonstrate similar zone of clearing in Δ*BCAL1086* and Δ*BCAL2974* compared to the K56-2 parent strain. C) Quantification of the zone of clearing demonstrates no significant alteration in siderophore activity in Δ*BCAL1086* and Δ*BCAL2974* compared to the K56-2 parent strain. D) Survival curve of *G. mellonella* infections. Data from three independent replicates of 8 to 10 larve for each biological group is shown with the standard deviation also denoted. Δ*BCAL1086* and Δ*BCAL2974* strains mirror the lethality of WT and Δ*pglL amrAB::native-pglL-his*_10_.

We also investigated whether the loss of either BCAL2974 or BCAL1086 drives proteome changes. Using label-free based quantitative proteomics we compared the proteomes of K56-2 WT, Δ*BCAL2974*, Δ*BCAL1086*, Δ*pglL*, Δ*cepR*, Δ*cepI* and Δ*pglL amrAB::S7-pglL-his*_10_ to assess the similarity between the proteomes as well as the specific proteins effected by the loss of these proteins. Proteomics analysis led to the identification of 3730 proteins with 2752 proteins quantified in at least 3 biological replicates in a single biological group (Supplementary Table 14). Clustering of the proteomic analysis revealed that Δ*BCAL2974* and Δ*BCAL1086* closely grouped with that of WT strains while, Δ*pglL*, Δ*cepR*, Δ*cepI* and Δ*pglL amrAB::S7-pglL-his*_10_ formed discrete clusters. This macro analysis indicated that mutations in BCAL2974 or BCAL1086 had a minimal effect on the proteome (Figure 7A, Supplementary Table 15A and B). Consistent with this conclusion, analysis of the specific proteins that varied between the different strains demonstrated few proteome alterations in Δ*BCAL2974* and Δ*BCAL1086* compared with Δ*pglL*, Δ*cepR* and Δ*cepI* (Figure 7B) with Δ*cepR*, Δ*cepI* and Δ*pglL* also demonstrating the expected similarility in their proteome changes (Fisher exact test Δ*cepR* vs Δ*pglL*, *p*-value 3.25 × 10^5^ and Δ*cepI* vs Δ*pglL*, 6.95 × 10^4^, Supplementary Table 16). Taken together the proteome analysis supports the contention that BCAL2974 and BCAL1086 have minimal effect on the proteome and are not responsible for the broad proteomic alterations observed in Δ*pglL*.

**Figure 7.**
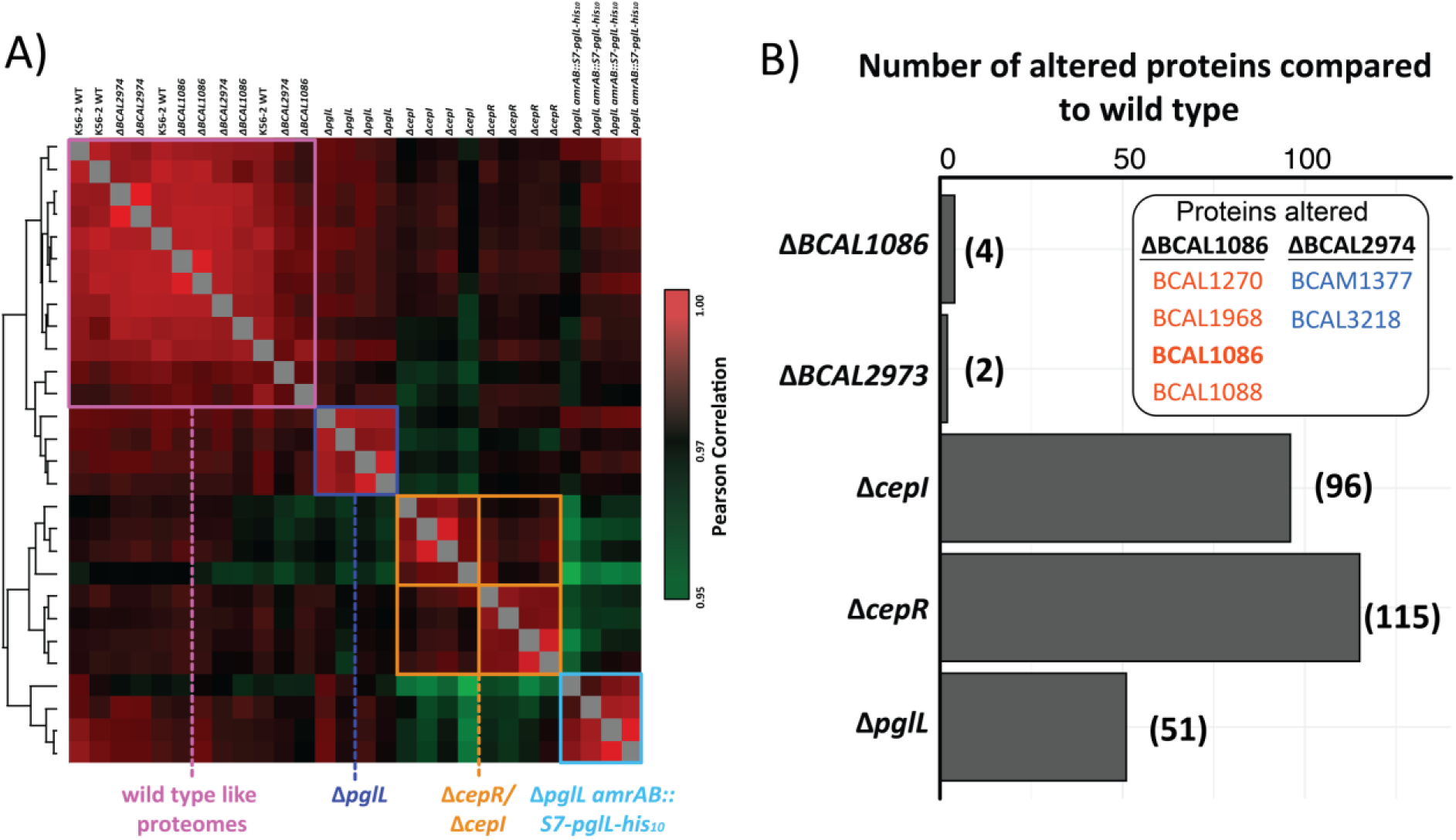
Disruption of BCAL1086 and BCAL2974 does affect the proteome as Δ*pglL*. A) Pearson correlation analysis of K56-2 WT, Δ*BCAL2974*, Δ*BCAL1086*, Δ*pglL*, Δ*cepR*, Δ*cepI* and Δ*pglL amrAB::S7-pglL-his*_10_ proteomes demonstrates K56-2 WT, Δ*BCAL2974*, Δ*BCAL1086* biological replicates cluster together while other strains form discrete clusters. B) Quantitative Proteome analysis of Δ*BCAL1086*, Δ*BCAL2974*, Δ*cepI*, Δ*cepR* and Δ*pglL* compared to wild type demonstrates minor proteome alterations compared to Δ*cepI*, Δ*cepR* and Δ*pglL*.

## Discussion

Although glycosylation is a common protein modification in bacterial species (49–51, 107) our understanding of how this modification influences bacterial physiology and pathogenesis is unclear. Recent insights into how glycosylation impacts bacterial proteomes have been obtained through study of the archetypical *N*-linked glycosylation system of *C. jejuni* (108, 109), yet it is unclear whether these observations are generalizable to another glycosylation systems such as *O*-linked glycosylation systems. Studies on the role of *N*-linked glycosylation within *C. jejuni* have revealed that defects associated with the loss of glycosylation stem from the loss of glycoproteins (108, 109) suggesting that *N*-linked glycosylation extends protein longevity in *C. jejuni*. In contrast, we find here that loss of *O*-linked glycosylation in *B. cenocepacia* has a more limited effect on the proteins targeted for glycosylation with only a subset of the known glycoproteins affected by the disruption of glycosylation (Figure 5). Therefore, the defect associated with loss of glycosylation in *B. cenocepacia* cannot be merely explained by protein instability. Indeed, we demonstrate that loss of glycosylation leads to changes in the expression of non-glycosylated proteins regulated by the CepR/I regulon (Figure 3), a global regulator of bacterial physiological changes including virulence and motility (39, 42, 48). Also consistent with a defect in the CepR/I regulon, loss of glycosylation results in lack of biofilm formation and reduced siderophore production. Therefore, our findings uncover a previously unknown link between loss of glycosylation and the repression of the CepR/I QS system.

The observation that biofilm formation is reduced in Δ*pglL* mirrors previous reports in *Acinetobacter baumannii* (55) and *C. jejuni* (63), but the link of this phenotype to alterations in regulations have not previously documented. Previous studies in *B. cenocepacia* have identified that not all CepR/I regulated proteins are required for biofilm formation. However, BapA (BCAM2143) plays a major role in the formation of biofilms on abiotic surfaces, whereas the lectin complex BclACB (BCAM0184–BCAM0186) contributes to biofilm structural development (45). Although BapA (BCAM2143) was not detected by our proteomic analysis, BclA and BclB (BCAM0186 and BCAM0184 respectively) were observed and decreased in Δ*pglL* (both -1.0 log_2_ decrease compared with WT, -log_10_(p): >3.05, Supplementary Table 7). Surprisingly, BclA and BclB increased in abundance in Δ*pglL*Δ*OGC* and Δ*OGC* strains (both 1.0 log_2_ increase compared with WT, -log_10_(p) >1.4, Supplementary Table 7), and these mutants formed extensive biofilms (Supplementary Figure 9). This result is consistent with recent work which has shown that with disruption of BCAL3116, the third gene in the OGC, resulted in enhanced biofilm formation (110) although the underlying mechanism driving this phenomenon is unclear. Concerning siderophore production, our proteomic data reveal that siderophore-associated proteins were reduced in both Δ*pglL* and Δ*OGC* (Figure 2) with glycosylation-null strains producing reduced zones of clearing in the CAS assays (Figure 4B and C, Supplementary Figure 10). However, the magnitude of the reduction in the CAS assays differed in the mutant since Δ*OGC* and Δ*pglL*Δ*OGC* presented significantly smaller zones of clearing than Δ*pglL* (Supplementary Figure 10). These results highlight that although the proteome changes observed in the glycosylation mutants Δ*pglL* and Δ*OGC* are highly similar they are not identical and show phenotypic differences. Therefore, a key question arising from our findings is how the loss of glycosylation alters gene regulation and whether the observed defects are simply the result of altered CepR transcriptional control. The lack of any glycosylated QS-associated proteins in *B. cenocepacia* (56) makes the identification of the link between a specific glycoprotein and QS changes unclear.

It is possible the observed alterations in biofilm formation and siderophore production are not solely driven by altered CepR regulation, but also reflect additional transcriptional alterations in the glycosylation null strains. This conclusion agrees with our observations of many differences in the abundance of transcriptional regulators in the DNA-associated proteome of glycosylation null strains (Figure 3C, Supplementary Table 10). An additional driver of these pleotropic effects may also be deleterious outcomes resulting from the manipulation of the *O*-linked glycosylation system. It has been suggested in *C. jejuni* that the disruption of glycosylation leads to undecaprenyl diphosphate decorated with *N*-linked glycan being sequestered from the general undecaprenyl diphosphate pool and that this depot effect may be a general phenomenon observed in all glycosylation mutants (64). Sequestration of undecaprenyl diphosphate was thought to drive an increase in the abundance of proteins within the non-mevalonate and undecaprenyl diphosphate biosynthesis pathways observed in glycosylation-null *C. jejuni* (64). However, in *B. cenocepacia* glycosylation mutants, we observe only minor alterations in the non-mevalonate (BCAL0802, BCAL1884, BCAL2015, BCAL2016, BCAL2085, BCAL2710, BCAM0911 and BCAM2738, Supplementary Figure 10A) and undecaprenyl diphosphate biosynthesis (BCAL2087 and BCAM2067, Supplementary Figure 10B) pathways, which argues against this phenomenon being common to all glycosylation mutants. Furthermore, the similarity of the proteome changes in the Δ*pglL*, Δ*OGC* and Δ*pglL*Δ*OGC* strains (Supplementary Figure 3) supports the conclusion that proteome changes are independent of the sequestration of the undecaprenyl diphosphate pool as Δ*OGC* and Δ*pglL*Δ*OGC* strains are unable build the *O*-linked glycan on undecaprenyl diphosphate.

Another explanation for the pleiotropic effects associated with loss of *O-*glycosylation could be the instability of the glycoproteins in the absence of the glycan. We identified two glycoproteins BCAL2974 and BCAL1086, both of unknown functions, which are reduced in abundance due to the loss of glycosylation. However, genetic experiments demonstrate that neither protein is responsible for the phenotypic and proteomic changes associated with loss of glycosylation (Figure 6 and 7). Further, in the case of BCAL1086, endogenous tagging and degradomic analysis confirm the loss of this protein in the Δ*pglL* background. Although these results support the breakdown of BCAL1086 as a consequence of the loss of glycosylation, an alternative explanation is that the changes in degradation arise from alterations in protease levels or activities in the Δ*pglL* mutant. Previously, we reported that Δ*pglL* results in enhanced casein proteolytic activity (65). However, our global proteome analysis shows only modest changes in protease levels. We also observed identical protease profiles from activity probe against multiple classes of protease in wild type, Δ*pglL* and Δ*pglL amrAB::S7-pglL-his*_10_ (Supplementary Figure 12), suggesting all of these strains have similar protease activities. More importantly, aside of glycoproteins BCAL2974 and BCAL1086, the other proteins targeted for glycosylation remain consistently stable in the glycosylation-defective mutants. Therefore, although loss of glycosylation may affect the stability of some glycoproteins, the pleiotropic effect found in the glycosylation mutants cannot be explained by alterations in protein degradation.

In summary, this work provides a global analysis of the impact of *O*-linked glycosylation on *B. cenocepacia* traits. The application of quantitative proteomics enabled the assessment of nearly half the predicted proteome of *B. cenocepacia* K56-2 and revealed a previously unknown link between *O*-linked glycosylation and the CepR/I regulon. The alteration in CepR transcription as well as its associated phenotypes support a model in which the virulence defects observed for glycosylation null strains arise from transcriptional changes and not from the direct result of glycosylation loss per se. This work challenges the idea that loss of glycosylation solely affects the stability and activity of the glycoproteome, and instead shows that glycosylation can influence the bacterial transcriptional profile and broader proteome.

## Acknowledgement

This work was supported by National Health and Medical Research Council of Australia (NHMRC) project grants awarded to NES (APP1100164) and Medical Research Council Confidence in Concept project CD1617-CIC04 (to MAV). NES was supported by an Overseas (Biomedical) Fellowship (APP1037373) and a University of Melbourne Early Career Researcher Grant Scheme (Proposal number 603107). LEM was supported by a Grimwade Fellowship from the Russell and Mab Grimwade Miegunyah Fund at The University of Melbourne and a DECRA Fellowship from the Australian Research Council (ARC, DE180100418). We thank the Melbourne Mass Spectrometry and Proteomics Facility of The Bio21 Molecular Science and Biotechnology Institute at The University of Melbourne for the support of mass spectrometry analysis. We also thank Silvia Cardona for kindly providing the plasmids pMS402, pCP300 and pPromCepR, Mario Feldman for pKM4, and the Canadian *Burkholderia cepacia* research and referral repository for providing K56-2. We would also like to thank David Thomas for his critical evaluation of the manuscript.

